# Intravenous BCG induces a more potent airway and lung immune response than intradermal BCG in SIV-infected macaques^1^

**DOI:** 10.1101/2024.07.17.603921

**Authors:** Solomon Jauro, Erica C. Larson, Janelle L. Gleim, Brendon M. Wahlberg, Mark A. Rodgers, Julia C. Chehab, Alondra E. Lopez-Velazques, Cassaundra L. Ameel, Jaime A. Tomko, Jennifer L. Sakal, Todd DeMarco, H. Jake Borish, Pauline Maiello, E. Lake Potter, Mario Roederer, Philana Ling Lin, JoAnne L. Flynn, Charles A. Scanga

## Abstract

Tuberculosis (TB), caused by *Mycobacterium tuberculosis* (Mtb), is one of the leading causes of death due to an infectious agent. Coinfection with HIV exacerbates Mtb infection outcomes in people living with HIV (PLWH). Bacillus Calmette-Guérin (BCG), the only approved TB vaccine, is effective in infants, but its efficacy in adolescents and adults is limited. Here, we investigated the immune responses elicited by BCG administered via intravenous (IV) or intradermal (ID) routes in Simian Immunodeficiency Virus (SIV)-infected Mauritian cynomolgus macaques (MCM) without the confounding effects of Mtb challenge. We assessed the impact of vaccination on T cell responses in the airway, blood, and tissues (lung, thoracic lymph nodes, and spleen), as well as the expression of cytokines, cytotoxic molecules, and key transcription factors. Our results showed that IV BCG induces a robust and sustained immune response, including tissue-resident memory T (T_RM_) cells in lungs, polyfunctional CD4+ and CD8αβ+ T cells expressing multiple cytokines, and CD8αβ+ T cells and NK cells expressing cytotoxic effectors in airways. We also detected higher levels of mycobacteria-specific IgG and IgM in the airways of IV BCG-vaccinated MCM. Although IV BCG vaccination resulted in an influx of T_RM_ cells in lungs of MCM with controlled SIV replication, MCM with high plasma SIV RNA (>10^5^ copies/mL) typically displayed reduced T cell responses, suggesting that uncontrolled SIV or HIV replication would have a detrimental effect on IV BCG-induced protection against Mtb.

## Introduction

Tuberculosis (TB) remains a major infectious cause of mortality globally. Over one-fourth of the world’s population is estimated to be or have been infected with *Mycobacterium tuberculosis* (Mtb), the causative agent of TB, and over 10 million new cases of active TB are reported yearly. In 2022, TB resulted in the death of 1.6 million people (1). TB disease is exacerbated in people living with HIV (PLWH), with morbidity rates 15 – 21 times higher than HIV-naïve individuals (2). PLWH have a higher likelihood of progression from latent to active TB as well as higher mortality rates, contributing to one out of three deaths in PLWH (1, 3). This grave effect of TB in PLWH correlates with the extent to which HIV infection depletes CD4+ T cells, an important cell type in host immunity against Mtb. CD4+ T cells are considered important for TB protection as HIV-mediated depletion of these cells results in rapid TB progression in PLWH (4, 5). However, even in some PLWH with normal CD4+ T cell levels (500-1500 cells/mm^3^), TB progression is more rapid than in HIV-naïve individuals (6). This may result from HIV-related factors that affect immunity, including impairment of anti-mycobacterial innate immunity by type I interferon (7, 8), immune suppression due to Treg/Th17 imbalances (9), interference of B cell signaling and memory formation (10), and inhibition of TNF-mediated macrophage activation and death (11, 12). A safe and effective TB vaccine would greatly reduce the risk of TB and mortality in PLWH and would significantly improve global health. Yet, HIV impairment of vaccine-elicited responses is not completely understood.

Bacillus Calmette-Guérin (BCG) is the only licensed vaccine for TB and was developed over a century ago. BCG is delivered intradermally (ID) to infants worldwide and is effective against severe TB in children, but its efficacy diminishes into adulthood (1). Many studies are focused on improving BCG efficacy. Clinical studies of BCG revaccination and protein subunit vaccines that examined sustained Quantiferon-TB conversion or prevention of latent TB progression to active TB demonstrated moderate vaccine efficacy of 30-50% in HIV-naïve subjects (13, 14). In nonhuman primates (NHP), BCG boosted with Mtb proteins improved T cell responses in the lungs but did not enhance protection from Mtb challenge (15). However, a study of BCG vaccination delivered by different routes in rhesus macaques revealed the intravenous (IV) route to be the most protective, conferring 90% protection from TB (16).

Despite these encouraging results, BCG, being a live attenuated vaccine, has the potential to cause disease in immunocompromised individuals. The risk of disseminated BCG in infants infected with HIV is significantly higher than in infants who are HIV-naïve (17), prompting safety concerns for BCG as a vaccine for immunocompromised persons (18). Nonetheless, we recently showed that IV BCG in macaques chronically infected with simian immunodeficiency virus (SIV), an NHP surrogate for HIV, was safe (at least with anti-BCG drug treatment post-vaccination), immunogenic, and protective (19).

Understanding vaccine-elicited immune responses that correlate with TB protection is critical for rational vaccine development. Several features have been associated with TB protection in IV BCG-vaccinated, SIV-naive macaques, including an influx of mycobacterial-specific T cells into the airways and expansion of T_RM_ cells in lung tissue (16, 20), elevation of mycobacterial-specific antibodies in circulation and in airway (21), and distinct transcriptional profiles in circulating PBMC (22). In a correlates analysis across a dose range of IV BCG, CD4 T cells expressing different combinations of IFNγ, TNF and/or IL-17 as well as the number of NK cells in airways were correlates of protection independent of BCG dose (20). In our study of IV BCG in SIV-infected macaques (16), TB protection tended to occur in animals with lower plasma viremia and higher levels of CD4+ T cells in the airways before Mtb challenge. Although we were able to assess vaccine responses in blood and airway, all animals underwent Mtb challenge, thereby complicating assessment of immune responses within tissues. Here, we evaluated immune responses to BCG in macaques chronically infected with SIV following vaccination delivered either IV or the standard ID route, and BCG responses in tissues were analysed without subsequent Mtb challenge. We used our well-characterized Mauritian cynomolgus macaque (MCM) model of SIV/Mtb coinfection (23–28) and infected the animals with SIV for 16 weeks before IV or ID BCG vaccination. Blood and airway cells were serially sampled, and animals were necropsied 5 months after vaccination to compare the immune response within various tissues between ID and IV BCG in SIV-infected MCM. We show that vaccination of SIV-infected macaques with IV BCG elicited an influx of CD4+, CD4+CD8α+, CD8αα+, CD8αβ+, and Vγ9+γδ, Vγ9-γδ T cells, and cytokine-producing CD4+ and CD8αβ+ T cells into the airways. Airway immune responses by IV BCG were mostly recapitulated in lung tissues with an expansion of CD8αα+, CD8αβ+, and γδ+ tissue-resident T cells.

## Materials and Methods

### Animals

Eleven (11) male Mauritius cynomolgus macaques (MCM) (*Macaca fascicularis*, 4-10 years old) were procured from Bioculture US (Immokalee, FL). Major histocompatibility complex (MHC) haplotypes, which were determined by MiSeq sequencing of PBMC at the Genomic Servies Core of the Wisconsin National Primate Research Center, and 8 the 11 MCM shared at least one copy of the M1 MHC haplotype. The MCM were screened for Mtb, as well as herpes B virus, SIV, SRV, and STLV, and housed at the University of Pittsburgh. All experimental procedures were conducted according to the ethical regulations of the University of Pittsburgh Institutional Animal Care and Use Committee (IACUC), and complied with the standards outlined in the Animal Welfare Act, the “Guide for the Care and Use of Laboratory Animals (8^th^ Edition)”, and “The Weatherall Report”. The University is fully accredited by AAALAC (accreditation number 000496), and its OLAW animal welfare assurance number is D16-00118. The IACUC reviewed and approved the study protocols 21048970 and 24044961. All veterinary procedures were performed under ketamine sedation. The MCM were monitored twice daily for any health issues.

### SIV infection

All MCM were infected intravenously with a 3×10^3^ tissue culture infectious dose (TCID_50_) of SIVmac239 which was grown in rhesus PBMC (Primate Assay Laboratory of the California National Primate Research Center, University of California, Davis CA). Plasma virus level was serially monitored by qPCR, as detailed below.

### BCG vaccination

Sixteen weeks after SIV infection, the MCM were randomly allocated into two vaccination groups: IV BCG (n=5), and ID BCG (n=6) (Figure 1a). Animals were vaccinated with BCG Danish Strain 1331, which was expanded at Colorado State University following the Aeras protocol (29). The BCG was prepared for injection by suspending in phosphate-buffered saline (PBS) containing 0.05% Tween 80 (Sigma-Aldrich) to prevent the inoculum from clumping. For ID inoculations, the upper arm was shaved and disinfected with chlorhexidine followed by isopropanol, and the standard human dose of 5×10^5^ CFU was injected into the dermis in a volume of 1 mL. For IV inoculation, the saphenous vein was used to inject 5×10^7^ CFU in 1 mL total volume, the dose we showed previously to confer high-order protection in SIV-infected MCM (17). To reduce the risk of BCG dissemination in these SIV-infected animals, the IV BCG-vaccinated MCM were treated with an anti-BCG drug regimen (isoniazid, 15mg/kg; rifampicin, 20mg/kg; ethambutol, 55mg/kg (HRE)) given by mouth daily for 8 weeks starting 3 weeks after vaccination, as done previously (17).

**Figure 1:**
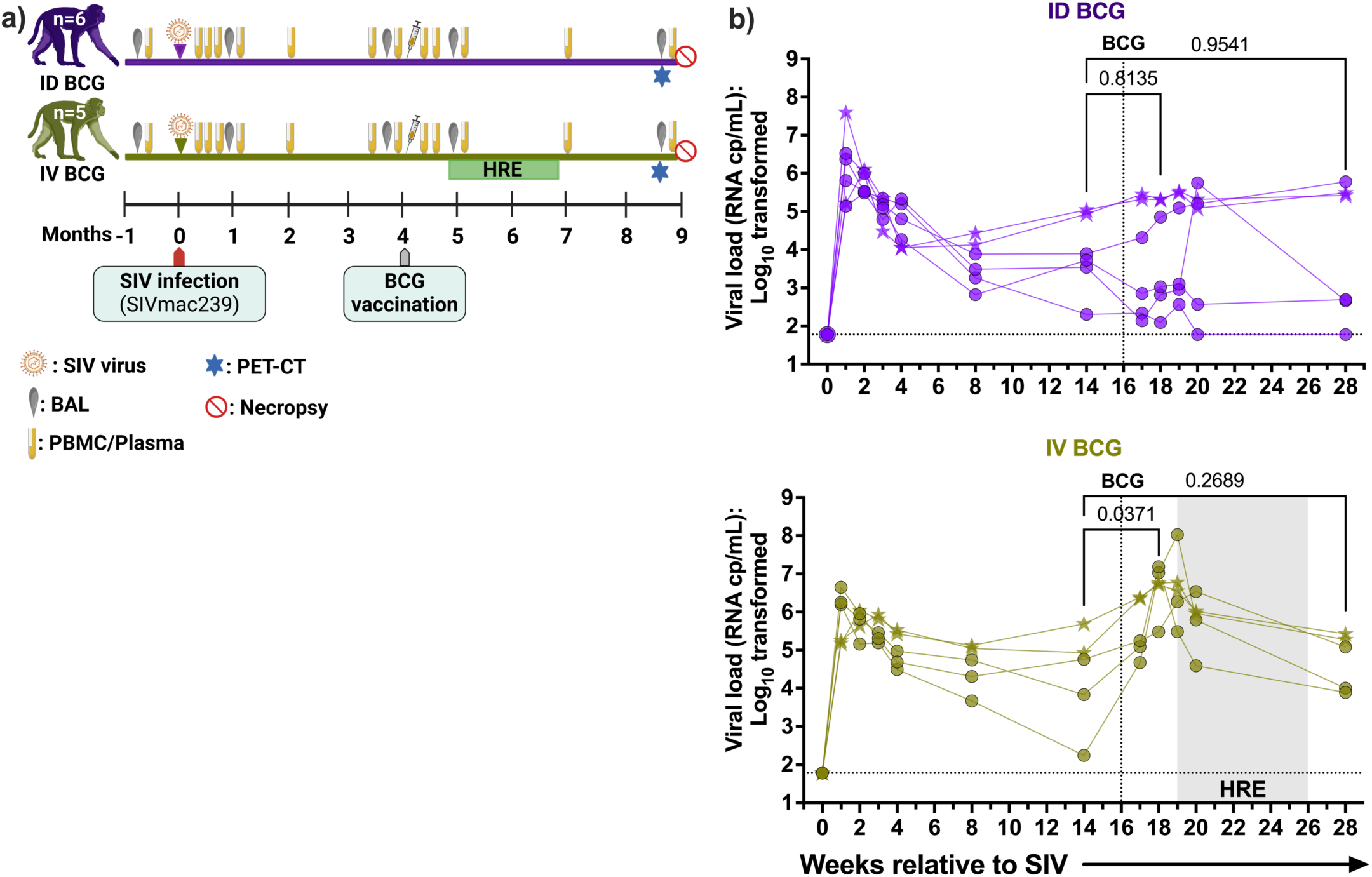
Study design and SIV plasma viremia. **a)** Study design indicating SIV infection, BCG vaccination, HRE treatment, serial BAL and blood collection, and PET-CT time points. **b)** Quantification of plasma viral copy equivalents over time in MCM vaccinated with ID BCG (top panel, purple; n=6) or IV BCG (bottom panel, green; n=5). The horizontal dashed line represents the LOQ of ∼62 copies/mL. Plasma viral load of each group was assessed by comparing two weeks before BCG vaccination to two- or 12-weeks after BCG vaccination using mixed-effects model with uncorrected Fisher’s LSD p-values reported. Each symbol represents an individual animal. SIV controllers and non-controllers (baseline >10^5^ copies/mL) are indicated by circles and stars, respectively.

### Sampling blood and airways

Whole blood was collected by venipuncture. PBMC were isolated using Ficoll-Paque PLUS gradient separation (GE Healthcare Biosciences) and cryopreserved in RPMI 1640/fetal bovine serum (FBS) containing 10% DMSO in liquid nitrogen. Bronchoalveolar lavage (BAL) was done by instilling and recovering warmed PBS (4 x 10 mL washes) under direct visualization using a pediatric bronchoscope. The recovered BAL fluid (BALF) was pelleted to collect the cells and a 15 mL aliquot of BALF was cryopreserved. The cells were resuspended into ELISpot media (RPMI 1640, 10% heat-inactivated human albumin, 1% L-glutamine, and 1% HEPES), counted, and apportioned for flow cytometric staining.

### PET-CT

PET-CT imaging using radiolabeled 2-deoxy-2-^18^Fluoro-D-glucose (FDG) was conducted within 3 days of necropsy. A MultiScan LFER-150 PET-CT scanner (Mediso Medical Imaging Systems, Budapest, Hungary) was used, as detailed previously (30, 31). Co-registered PET-CT images were analyzed using OsiriX MD software (v 12.5.2, Pixmeo, Geneva, Switzerland) to measure the total FDG avidity of the lungs, thoracic lymph nodes (32). Total FDG avidity correlates with overall inflammation.

### IV CD45 staining for tissue-resident cells

To differentiate T_RM_ cells from intravascular cells, IV anti-CD45 antibody staining was performed as previously described (16, 33, 34). Briefly, 5 minutes before euthanasia, animals were maximally bled and then slowly infused intravenously with 5 mL of purified Alexa Fluor 488-conjugated αCD45 antibody to achieve a dose of 100 μg/kg. For the ID BCG vaccination group, 5 of 6 macaques were injected IV with anti-CD45 antibodies. When harvested tissues were analyzed by flow cytometry, CD45 immune cells present in parenchyma were unstained, while those in the vascular were positively stained with anti-CD45 antibody.

### Necropsy and tissue processing

Five months after vaccination, animals were humanely euthanized and necropsied to collect lung tissue, thoracic and peripheral lymph nodes, liver, and spleen. Each tissue sample was divided and a portion was fixed in neutral-buffered formalin for histopathology, with the remainder homogenized to yield a single cell suspension, as previously described (24). To detect any viable BCG, serial dilutions of this homogenate were plated onto 7H11 agar and incubated at 37°C, 5% CO_2_ for three weeks. Single cell suspensions were portioned immediately for flow cytometric staining with the remainder cryopreserved in RPMI 1640/FBS containing 10% DMSO in liquid nitrogen. Formalin-fixed tissue was embedded in paraffin, sectioned, and stained with hematoxylin and eosin for microscopic examination.

### SIV plasma viremia

The number of SIV RNA copies per milliliter was measured using a two-step real-time qPCR assay. Viral RNA was extracted from 500 µL of plasma using an automated sample preparation platform QIAsymphony SP (Qiagen, Hilden, Germany), along with a Virus/Pathogen DSP midi kit and the *cellfree500* protocol (Qiagen). The extracted RNA was annealed to a reverse primer specific to the gag gene of SIVmac239 (5’-CAC TAG GTG TCT CTG CAC TAT CTG TTT TG −3’), then reverse transcribed to obtain cDNA using SuperScript™ III Reverse Transcriptase (Thermo Fisher Scientific, Waltham, MA) along with RNAse Out. The resulting cDNA was treated with RNAse H and combined with a custom 4x TaqMan™ Gene Expression Master Mix (Thermo Fisher Scientific) containing primers and a fluorescently labeled hydrolysis probe specific for the gag gene of SIVmac239 (forward primer 5’-GTC TGC GTC ATC TGG TGC ATT C −3’, reverse primer 5’-CAC TAG GTG TCT CTG CAC TAT CTG TTT TG −3’, probe 5’- /56-FAM/CTT CCT CAG TGT GTT TCA CTT TCT CTT CTG CG/3BHQ_1/ −3’). The qPCR was performed using a StepOnePlus™ or QuantStudio 3 Real-Time PCR System (Thermo Fisher Scientific). To determine the SIV gag RNA copies per reaction, quantification cycle data and a serial dilution of a well-characterized custom RNA transcript containing a 730 bp sequence of the SIV gag gene were used. The mean RNA copies per mL were calculated by adjusting for the dilution factor. The limit of quantification (LOQ) was determined based on the exact binomial confidence boundaries and the percent detection of the target RNA concentration, which is approximately 62 copies per mL.

### Multiparameter flow cytometry

Cells from BAL, PBMC, and tissue collected at necropsy were stained for flow cytometric analysis. BAL and necropsy tissue were stained promptly upon collection, while cryopreserved PBMC were stained as a batch. Cells were seeded at 1 x 10^6^ cells/well in a 96-well plate in ELISpot media, then stimulated with either Mtb H37Rv whole cell lysate (WCL, 20 μL/mL; BEI Resources) or PDBu and ionomycin (P&I) for 6 hours at 37°C. After 2 hours of incubation, Golgi Plug (BD Biosciences, Cat no. 555029) was added at 8 μg/mL, incubated for an additional 4 hours, and washed twice with PBS. Live/Dead stain was added and incubated for 10 minutes at room temperature in the dark, then washed twice with PBS. A cocktail of the surface stain (Supplemental Table 1) supplemented with BD Horizon™ Brilliant Stain buffer (BD Biosciences, Cat no. 566349) was added to each well (50 μL/well). After 20 minutes of incubation at 4°C, the cells were washed twice and permeabilized for intracellular staining (ICS) with BD Cytofix/Cytoperm™ (BD Biosciences, Cat no. 554722) for 10 minutes at room temperature. For intranuclear staining (INS) of transcription factors (necropsy samples only), 1x True-Nuclear™ (BD Biosciences, Cat. No. 424401) was added and incubated for 45 minutes. The cells were then washed twice with 1x perm wash for ICS or True-Nuclear™ 1x Fix Perm wash buffer for INS. Staining was done with 50 μL/well of the ICS and INS antibody cocktails (Supplemental Table 1) for 20 and 30 minutes, respectively. This was followed by washing twice and resuspending the stained cells in 150 μL of FACs buffer. Samples were run on a BD Cytek Aurora and analyzed using FlowJo Software for Mac OS (v10.8.1). Detailed gating strategies are shown in Supplemental Figure 3a-d.

### Plasma and BAL antibody ELISA

Plasma and 10-12 fold concentrated BALF were used to measure IgG and IgM titers to Mtb H37Rv WCL and lipoarabinomannan (LAM), as previously described (16). Briefly, 1 ug of WCL and LAM were coated onto 96-well Costar^®^ Assay plate (Nunc). Coated plates were blocked with PBS/10% FBS for two hours at room temperature and washed with PBS/0.05% Tween 20. Concentrated BALF or plasma 1:5 serially diluted (8 dilutions/sample) were then incubated for two hours at 37 °C and washed. Then 100 uL/well of HRP-conjugated goat anti-monkey IgG h+l (50 ng/ml; Bethyl Laboratories, Inc.), or IgM chain (0.1 ug/ml; Rockland Immunochemicals Inc.) was added and the plates incubated for 2 hours at room temperature. Plates were developed with 100 uL/well Ultra TMB^®^ substrate (Invitrogen) and fixed with 100 uL/well 2N sulfuric acid. Plates were read at 450 nm using a Spectramax190 microplate reader (Molecular Devices) and presented as endpoint titers for IgG or midpoint titers for IgM when samples did not titer to a cut-off. Endpoint titers are reported as the reciprocal of the last dilution with an optical density (OD) above the detection limit or twice the OD of an empty well.

### Statistical analysis

All the data were transformed as log_10_(x + 1). Longitudinal BAL, PBMC, and SIV viremia data were analyzed using a mixed model fit using the Restricted Maximum Likelihood (REML) with the Geisser-Greenhouse correction. Individual animals were considered a random variable. Fixed effect tests were used to evaluate the differences between different time points, vaccine route, or vaccine route versus time points. All results from pairwise comparisons of interest (comparisons among time points within vaccine group and comparison between IV BCG vs ID BCG within time points) were reported as uncorrected Fisher’s Least Significant Difference (LSD) p-values. Shapiro-Wilk test was used to test data for normality. Comparisons between IV BCG and ID BCG were made using the unpaired t-test for normally distributed data or the Mann-Whitney test for nonparametric data. Bivariate relationships were assessed using Pearson’s correlation coefficient (r). All tests conducted in the study were two-sided, and statistical significance was determined at a significance level of *P* < 0.05. *P-*values ranging from 0.05 to 0.10 were deemed indicative of a trend. GraphPad Prism 10 for MacOS (Version 10.0.2) was used for statistical analyses.

## Results

### IV BCG transiently increases plasma SIV viremia

The study was designed to compare immune responses to IV or ID BCG in SIV-infected MCM without the confounding variable of Mtb challenge. Animals were infected with 3,000 TCID_50_ SIVmac239 for 16 weeks and then randomized into two groups to be vaccinated with BCG by either the IV route (n=5) or the ID (n=6) route (Figure 1a). Animals in the IV BCG group were treated for two months with HRE, an anti-BCG drug combination, starting 3 weeks after vaccination to reduce the risk of BCGosis in these immunocompromised animals (19). Plasma viremia peaked within 2 weeks of SIV infection, then spontaneously fell to the viral set-point unique to each animal (Figure 1b). Animals with plasma viremia set points above 10^5^ copies/ml were considered to be “non-controllers”, indicating an inability to naturally control SIV replication (35–37). Two animals in each vaccination group were non-controllers by 14 weeks post-SIV infection (pre-BCG) (Figure 1b, non-controllers marked with star symbols). The median viremia at 14 weeks (pre-vaccination) was higher in the IV BCG group (Figure 1b, bottom panel). ID BCG vaccination did not significantly affect plasma viremia (Figure 1b, top panel), although SIV replication transiently increased in one animal 3 weeks after ID BCG. In contrast, IV BCG elicited a significant increase in plasma viremia in all animals within 2 weeks, which spontaneously returned to the approximate pre-BCG setpoints in most animals over the next 10 weeks (Figure 1b, bottom panel), as seen previously (19).

### IV BCG elicits more T cells in the airway

Immune responses in the airway are associated with IV BCG-elicited protection against Mtb infection (16, 34). Analysis of BAL samples taken 4 and 15 weeks after SIV infection (before vaccination) revealed no significant difference in total leukocyte and T cell counts from pre-SIV levels (Figure 2a, b). However, CD4+ T cells were significantly reduced and CD4+CD8α+ T cells were modestly reduced 4 weeks after SIV infection in both groups (Figure 2b). Both cell populations returned to pre-SIV levels prior to BCG vaccination, except in animals with high viremia (Figure 2b). B cells and CD11b-CD11c+ (likely dendritic cells) were also significantly depleted in the ID BCG groups following SIV infection (Supplemental Figure 1a).

**Figure 2:**
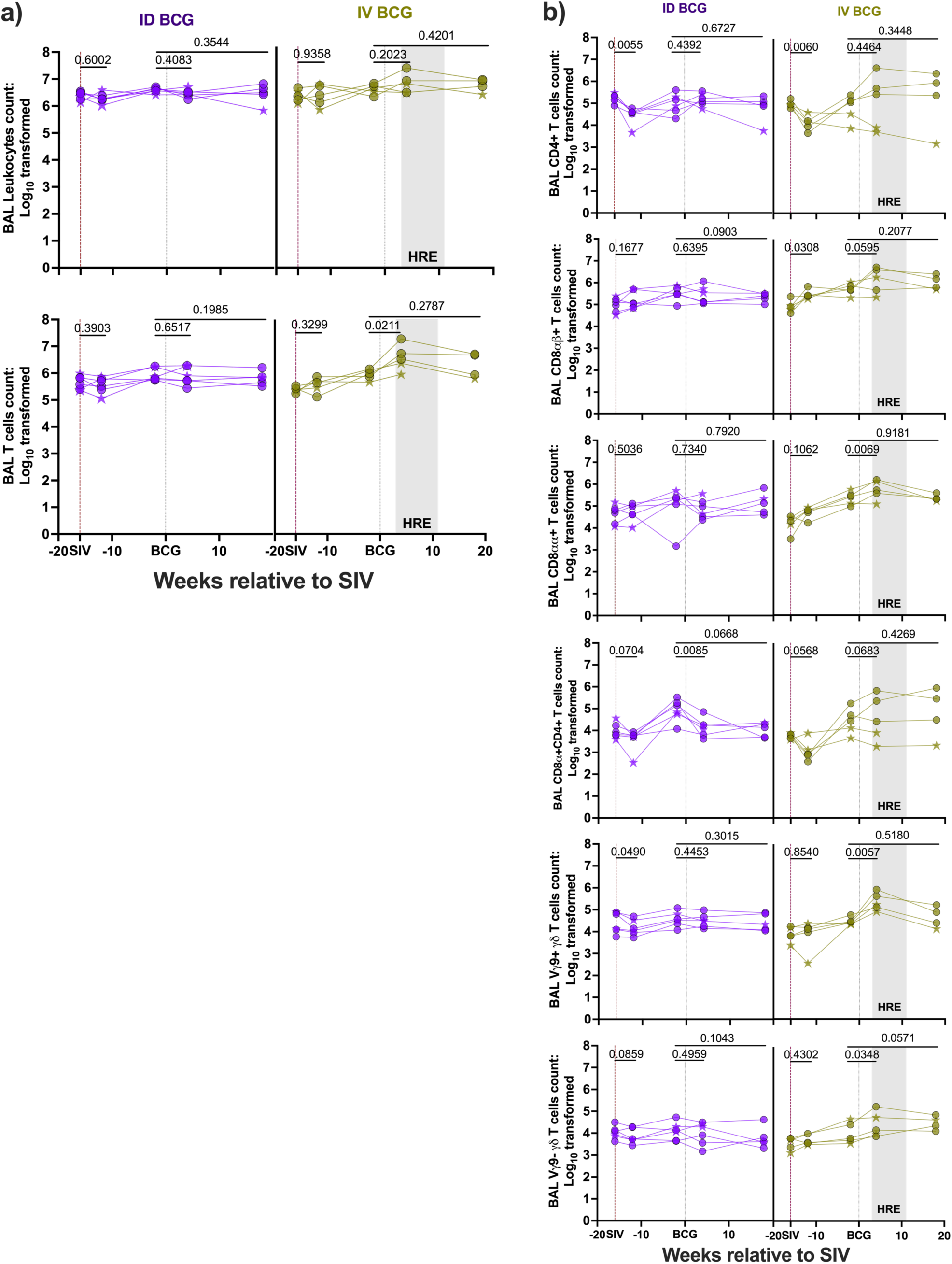
Immune cell composition in airways during SIV infection and BCG vaccination. **a)** Numbers of total leukocytes and T cells in BAL. **b)** Numbers of T cell subsets in BAL. Each point represents an individual BAL with SIV non-controllers indicated by stars. Mixed-effects model with uncorrected Fisher’s LSD p-values (Table 1) are shown.

**Table 1:**
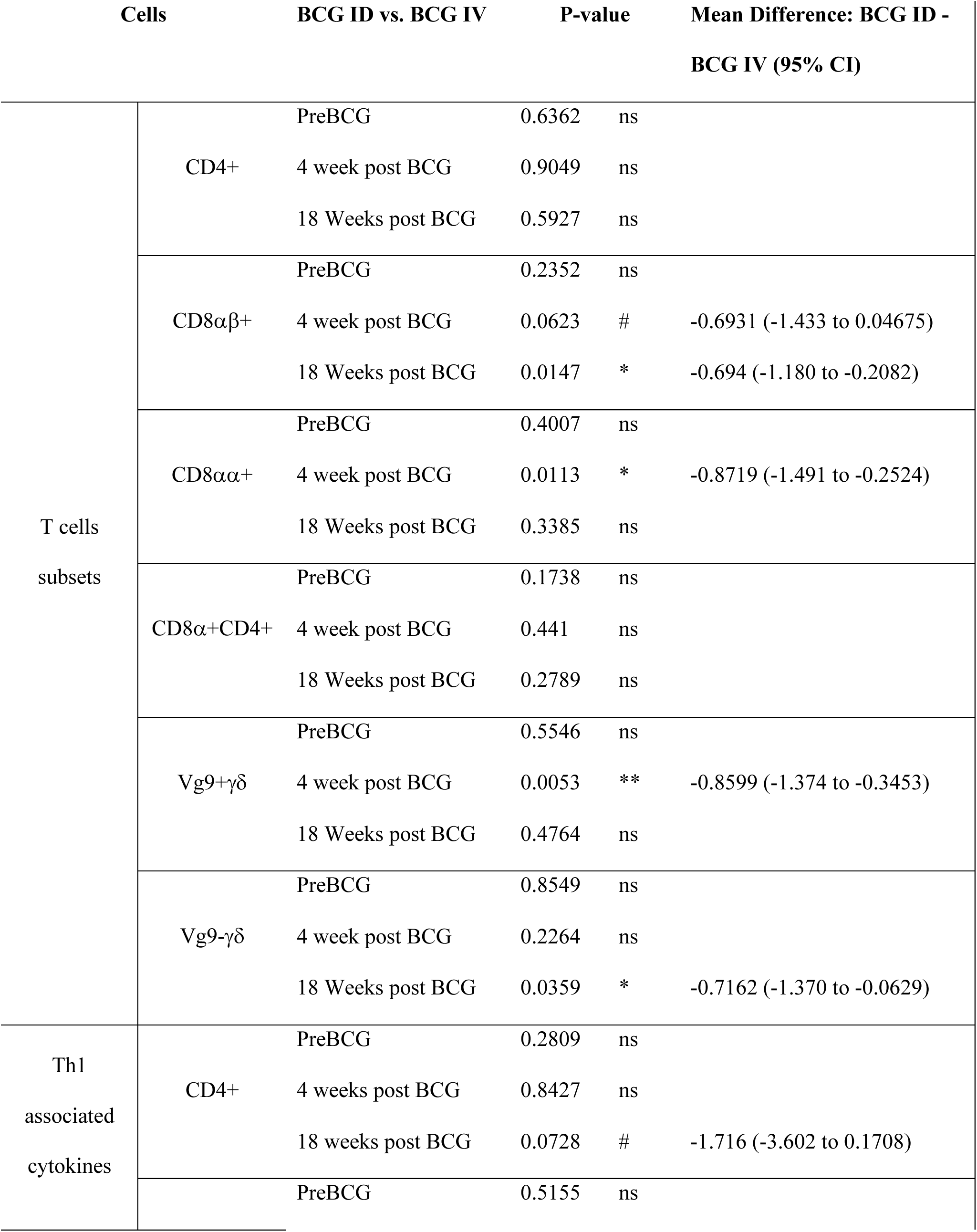
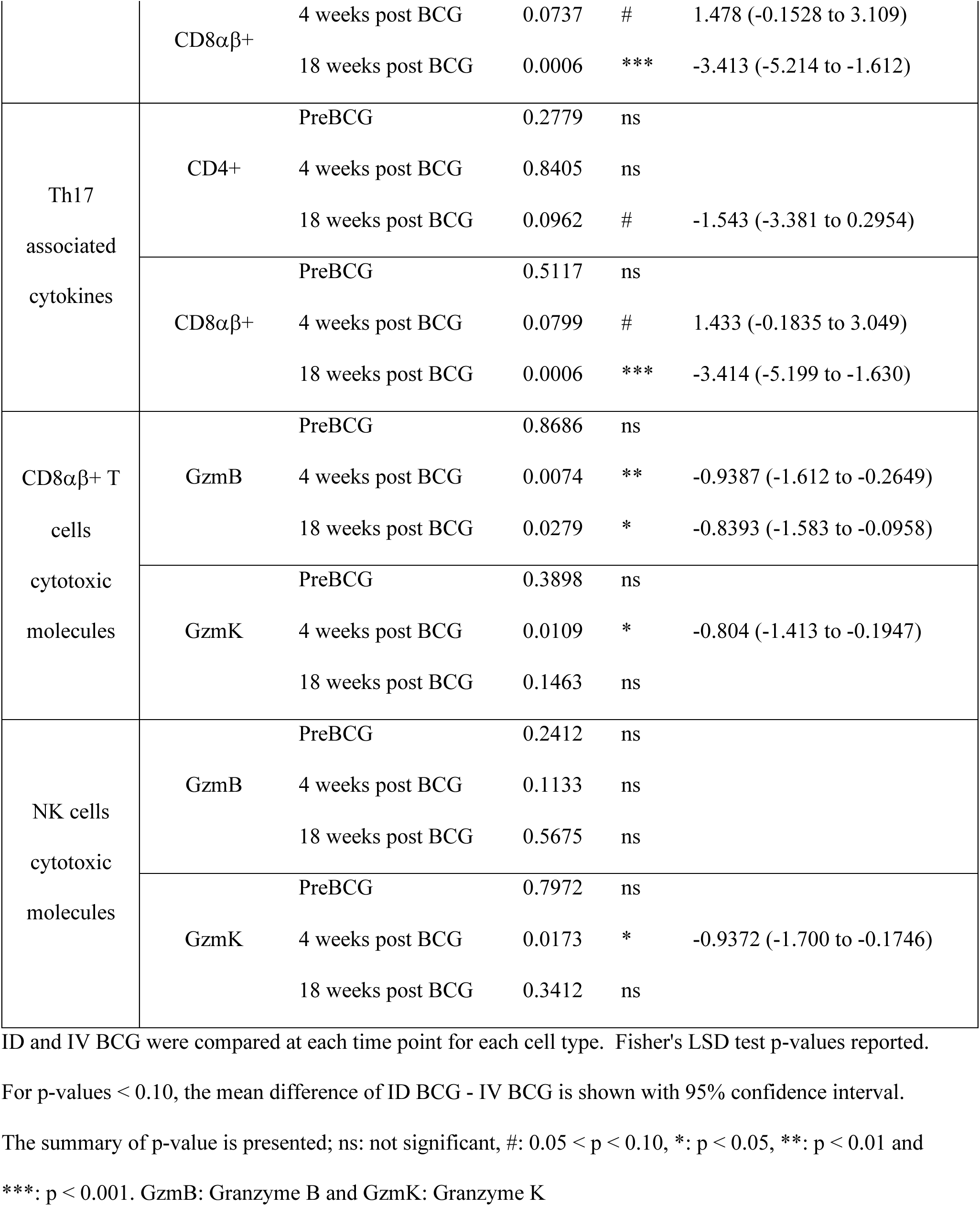
Mixed model analysis comparing cellular immune response elicited by ID BCG versus IV BCG vaccination routes at different time points.

Following IV BCG vaccination, animals exhibited modest increases in leukocyte count and significant increases in T, B, and NK cells in the airways. These cell counts remained unchanged in the ID-vaccinated animals (Figure 2a, Supplemental Figure 1a). Notably, IV BCG, but not ID BCG, induced significant increases in CD8αβ+, CD8αα+, Vγ9+γδ, and Vγ9-γδ T cells and modest increases in CD4+, and CD4+CD8α+ T cells 4 weeks after vaccination (Figure 2b, and Table 1). These cell types returned to pre-BCG levels by 18 weeks post-vaccination, except Vγ9-γδ T cells which remained elevated (Figure 2a, 2b, and Table 1). There was no correlation between airway CD4+ T cell levels with SIV viremia in either group (Data not shown). Thus, we show IV BCG but not ID BCG elicits an influx of T cells into the airway of SIV-infected MCM.

### IV BCG elicits more robust airway T cell responses than ID BCG

We evaluated the ability of CD4+ and CD8αβ+ T cells in the airways to produce cytokines, focusing on those involved in T1 (IFNγ, IL-2, TNF) and T17 (IFNγ, IL-2, IL-17, TNF) responses due to their important roles in TB protection (20, 38, 39). Cytokine responses were assessed after *ex vivo* restimulation with Mtb WCL to measure mycobacterial-specific responses. The number of antigen-specific CD4+ T cells in the airways producing T1- and T17-related cytokines significantly increased 4 weeks after BCG, regardless of vaccine route (Figure 3a). T1 CD4+ T cell responses remained significantly elevated up to 18 weeks in only the IV BCG group, while T17 responses remained elevated in both IV- and ID-vaccinated groups (Figure 3a). In contrast, only ID BCG elicited significant T1 and T17 responses in airway CD8αβ+ T cells by 4 weeks post-vaccination. However, by 18 weeks after vaccination, T1 and T17 CD8αβ+ T cells were significantly increased in IV-vaccinated animals, while those responses were waning in the ID-vaccinated animals (Figure 3b). In summary, BCG vaccination by both the ID and IV routes elicited cytokine responses in airway CD4+ T cells by 4 weeks, but CD8αβ+ T cell responses were delayed following IV BCG. IV BCG resulted in more sustained CD4+ and CD8αβ+ T cell responses than ID BCG.

**Figure 3:**
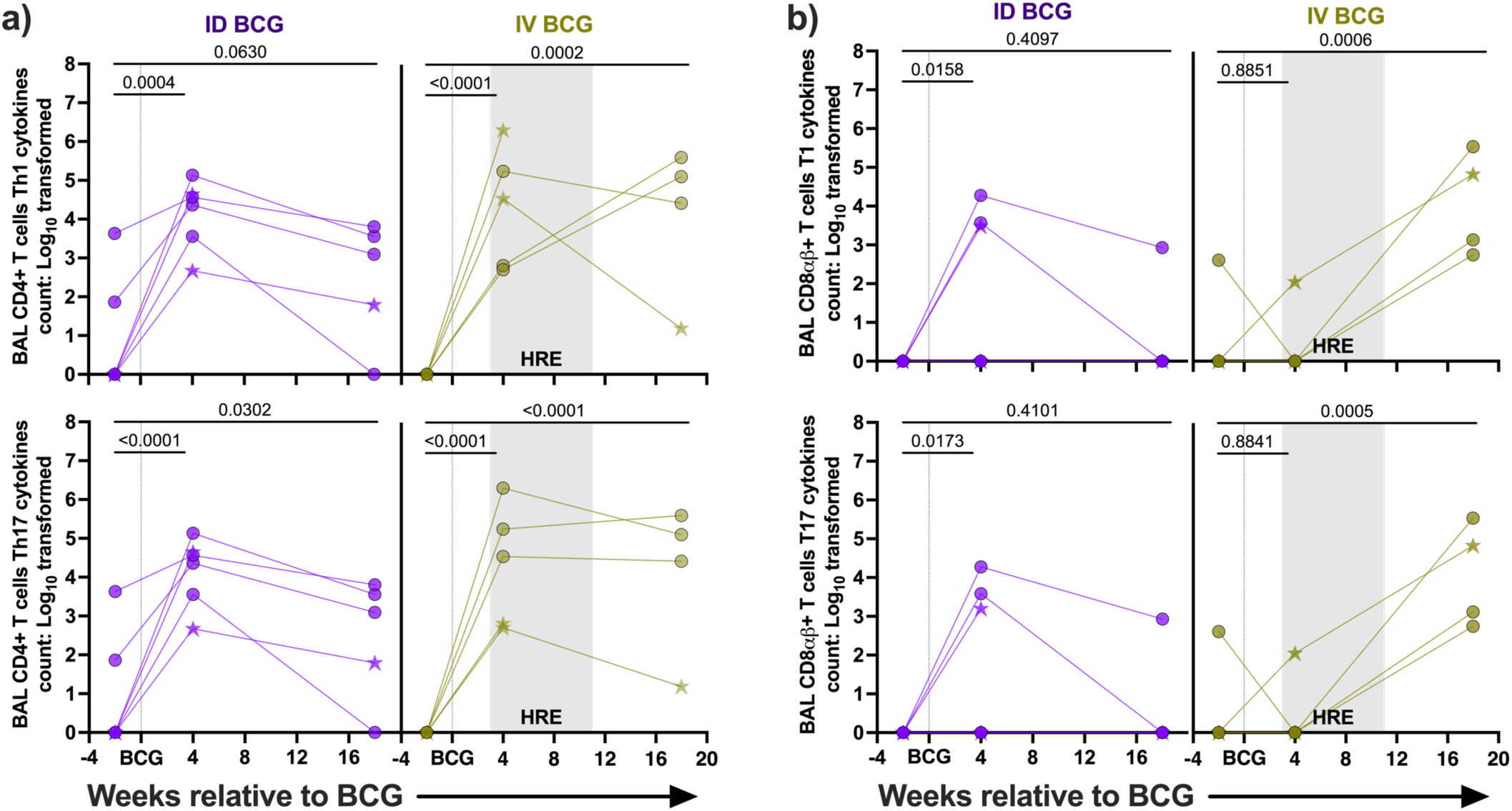
T cell cytokines in airways following BCG vaccination. **a)** The number of CD4+ T cells making mycobacterial-specific T1 and T17 responses in airways. **b)** The number of CD8αβ+ T cells making mycobacterial-specific T1 and T17 responses in airways. SIV non-controllers are indicated by stars. Mixed-effects model with uncorrected Fisher’s LSD p-values between groups are reported in Table 1.

### Only IV BCG induces an influx of cytotoxic CD8αβ+ and NK cells in airways

We next evaluated production of the cytotoxic effector granzyme B (GzmB) and granzyme K (GzmK) by CD8αβ+ T cells and NK cells recovered from the airways. Cytotoxic effector may contribute to Mtb protection by lysing infected cells and/or intracellular killing of bacteria (40–42). SIV infection significantly increased both GzmB+ and GzmK+ CD8αβ+ T cells in both groups (Figure 4a), but did not affect the number of NK cells producing granzymes (Figure 4b). Following vaccination, IV BCG, but not ID BCG, elicited an additional and significant boost in CD8αβ+ T cells producing GzmB or GzmK, with GzmB+ cells remaining modestly elevated 18 weeks after vaccination. IV BCG also significantly increased the number of NK cells producing either GzmB or GzmK, but returned to pre-vaccination levels 18 weeks after BCG (Figure 4b and Table 1). These data demonstrate that IV BCG induces an early influx of cytotoxic effectors from CD8αβ+ T cells and NK in the airways.

**Figure 4:**
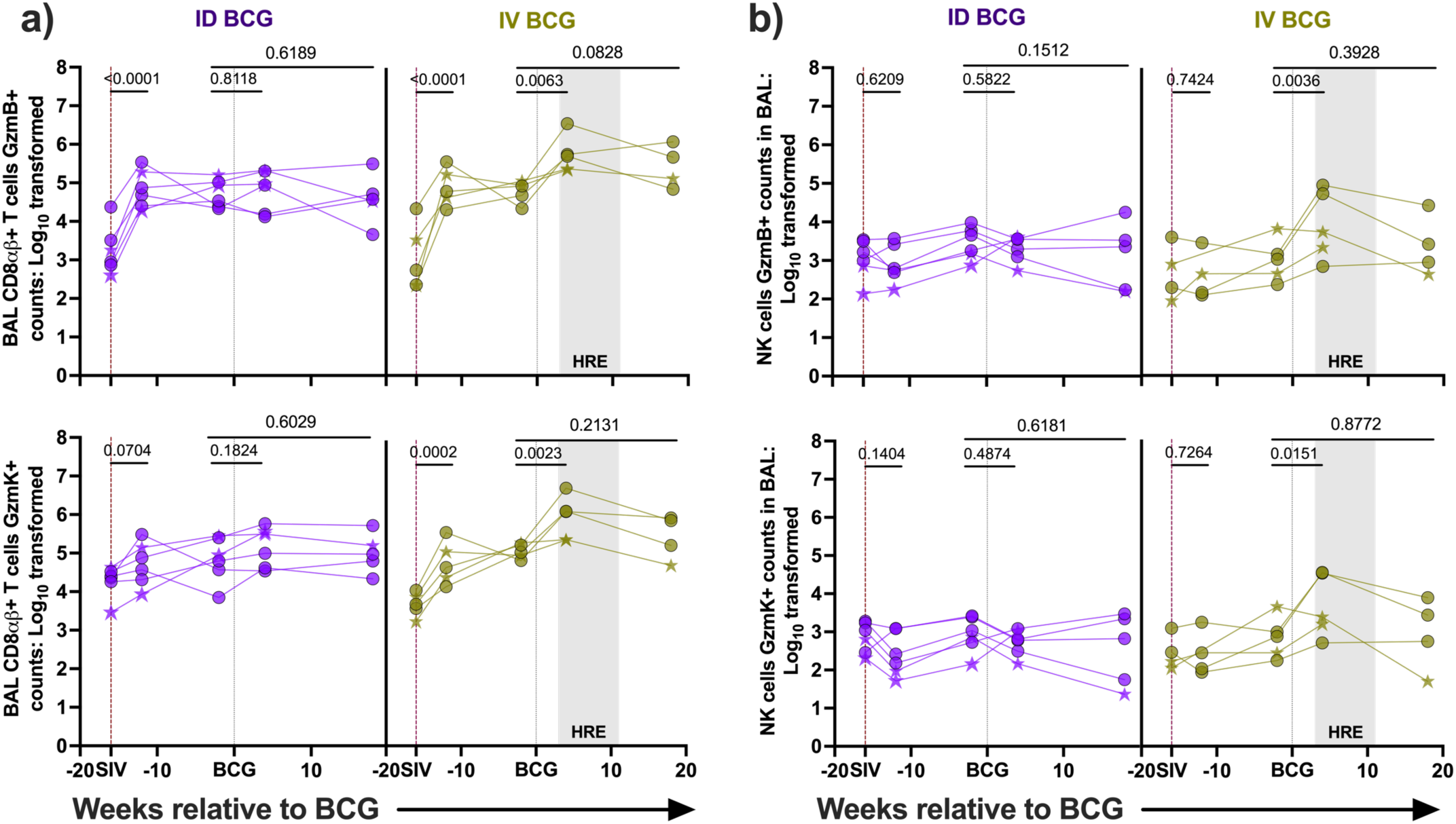
CD8αβ+ T cells and NK cells cytotoxic effector in airways. **a)** The number of CD8αβ+ T cells producing GzmB or GzmK. **b)** The number of NK cells producing GzmB or GzmK. SIV non-controllers are indicated by stars. Mixed-effects model with uncorrected Fisher’s LSD p-values between groups are shown Table 1.

### Transient changes in circulating immune populations following vaccination

PBMC were analyzed by flow cytometry to assess effects of BCG vaccination on circulating cell populations. Following SIV infection, the number of CD4+ and Vγ9+γδ T cells were significantly depleted in the IV BCG and ID BCG groups, respectively, prior to vaccination (Supplemental Figure 1c). Surprisingly, IV BCG vaccination induced a significant depletion in the number of NK, B cells, CD4+ and CD4+CD8α+ T cells by 4 weeks post-vaccination, while Vγ9+γδ T cells were depleted 4 weeks after ID BCG-vaccination. These cell types returned to their pre-BCG levels by 18 weeks post-vaccination, although a significant decrease in NK cells remained at this late time point in ID-vaccinated animals (Supplemental Figure 1b, 1c).

### IV BCG induces mycobacterial-specific antibody response in plasma and airways

We used ELISAs to measure WCL- and LAM-specific antibodies in plasma and BALF after BCG vaccination. IgG and IgM levels were associated with Mtb protection in IV BCG-vaccinated macaques (16, 19, 21, 43). Here, we observed a significant increase in plasma IgG specific for WCL and LAM following both ID and IV BCG by 4 weeks (Figure 5a, top panels). However, LAM-specific IgG in plasma returned to pre-BCG levels in both IV and ID BCG macaques by 18 weeks post-vaccination, while WCL-specific IgG returned to pre-BCG levels only in the ID BCG group (Figure 5a). Both IV and ID BCG elicited significant WCL-specific IgM in plasma, but only IV BCG induced significant LAM-specific IgM 4 weeks post-vaccination (Figure 5a, bottom panels). By 18 weeks after BCG, the IgM responses remained significant in the IV-compared to the ID-vaccinated group (Figure 5a, Table 2).

**Figure 5:**
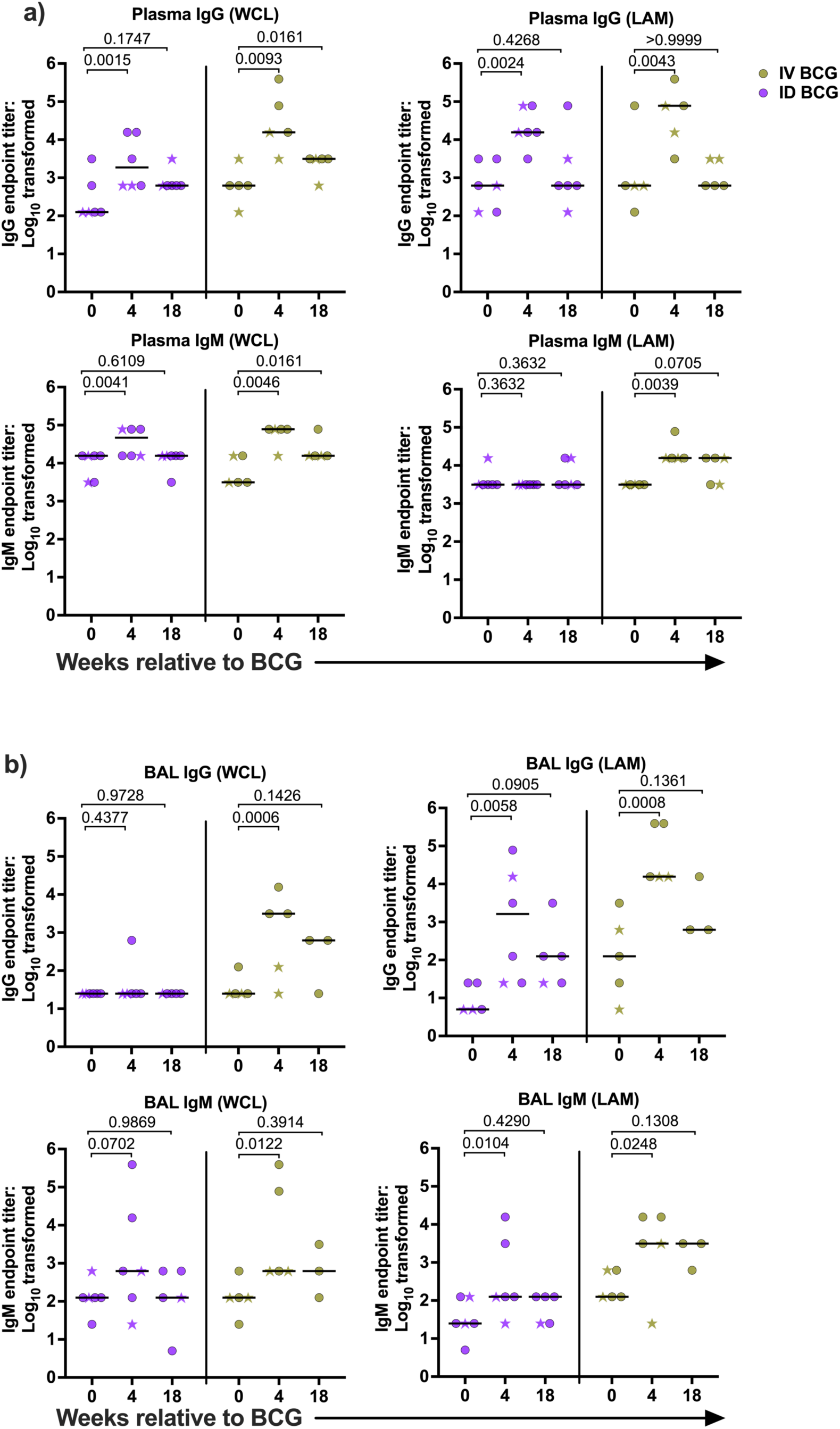
Mycobacteria-specific antibodies in BAL and plasma following BCG vaccination. IgG and IgM that bind WCL or LAM in plasma **(a)** and BALF **(b)**. SIV non-controllers are indicated by stars. Mixed-effects model with uncorrected Fisher’s LSD p-values in Table 2 represents group comparison.

**Table 2:**
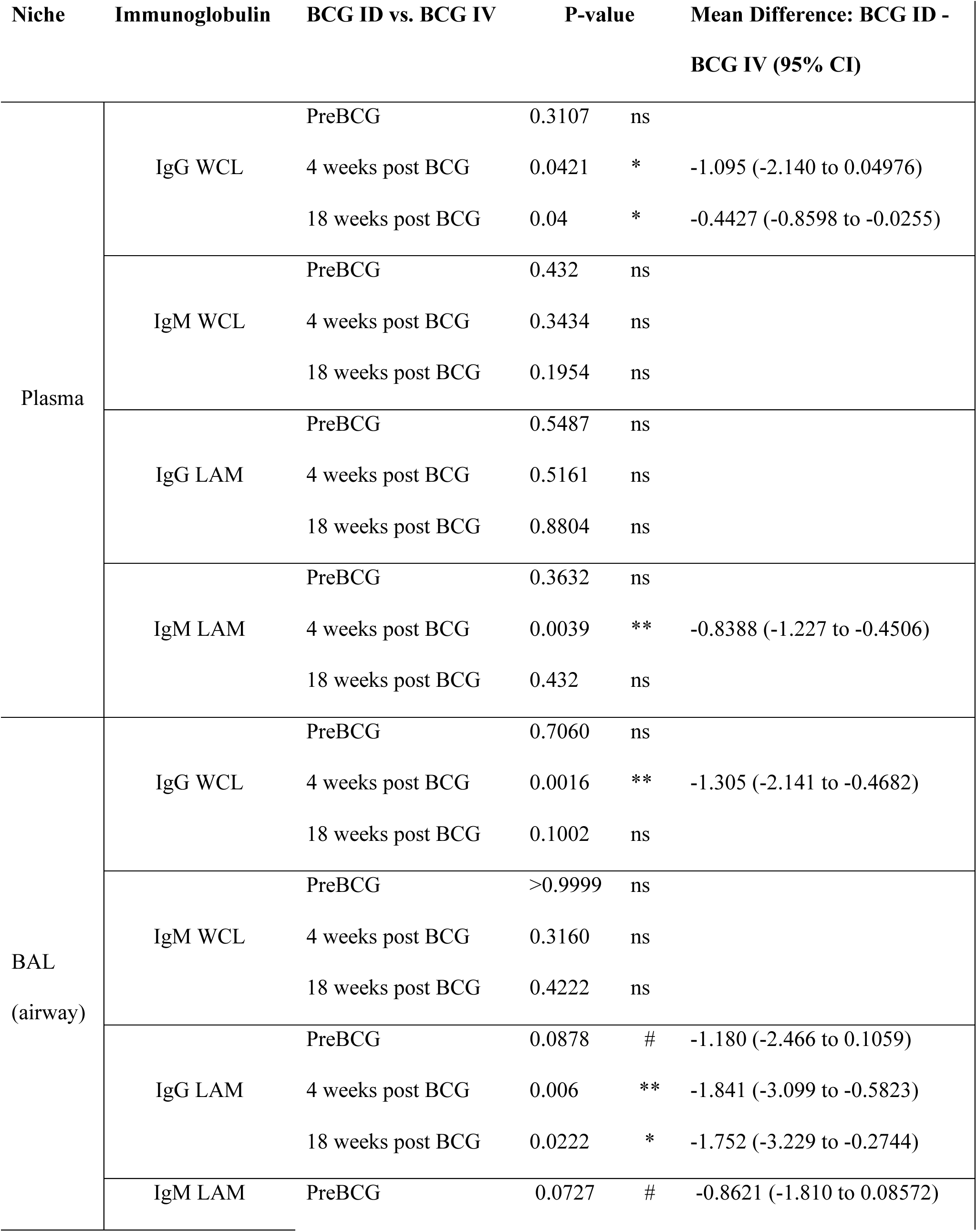
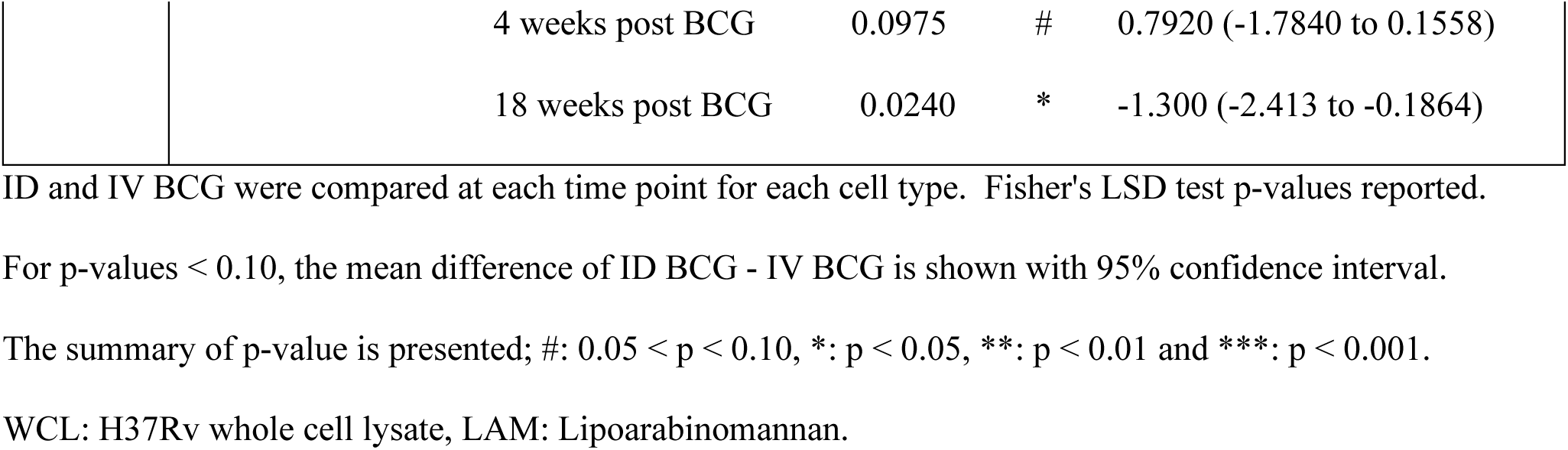
Mixed model analysis comparing the humoral immune response elicited in the plasma and airways by ID BCG versus IV BCG at different time points.

In airways, IV BCG, but not ID BCG, elicited a significant increase in WCL-specific IgG and IgM while LAM-specific IgG and IgM responses were significant in both IV and ID BCG 4 weeks after vaccination (Figure 5b). However, both WCL- and LAM-specific antibodies declined to pre-BCG levels by 18 weeks after vaccination. (Figure 5b). Comparing each time point between IV and ID BCG showed that IV BCG elicited higher levels of LAM- and WCL-specific IgG in the airways at 4 weeks post-BCG than ID BCG. However, by 18 weeks after BCG vaccination, only LAM-specific IgG and IgM were significant in the airway of IV BCG compared to ID BCG (Table 2).

### IV BCG induces robust immune responses in lung tissue

We interrogated immune responses across various tissues in BCG-vaccinated SIV+ MCM without the confounding variable of Mtb challenge, something that was not possible in our previous study in which all SIV+, BCG-vaccinated animals underwent Mtb challenge (19). Here, we compared T cell responses in lungs, thoracic lymph nodes, and spleen five months after vaccination. Since T_RM_ cells were associated with IV BCG-elicited protection (16), we included anti-CD69 antibody, a canonical marker of T_RM_ cells and activation (33), in our flow panels (Supplemental Table 1). To differentiate T_RM_ immune cells from those in circulation, fluorescently-labeled anti-CD45 antibody was administered just prior to euthanasia (16, 34). This analysis revealed significantly more T_RM_ cells in lungs of animals that received IV BCG compared to ID BCG (Figure 6a). Of the other T_RM_ cell subsets identified, there were significantly more CD8αβ+, CD8αα+, and γδ+ T cells in lung tissue of IV BCG-vaccinated animals (Figure 6b). CD4+ and CD4+CD8α+ T cell numbers did not differ significantly between the groups, most likely due to the low CD4+ T cell numbers in the MCM with higher SIV viral levels (Figure 6b). Indeed, CD4+ T cells in lung were modestly inversely correlated with SIV plasma viremia in both ID and IV BCG MCM (Data not shown). T_RM_ cell subset numbers in thoracic lymph node were similar in animals vaccinated with either ID or IV BCG (Supplemental Figure 2a), and the number of CD4+ and CD8α+CD4+ T cells in thoracic lymph nodes appeared independent of SIV viremia level (Supplemental Figure 2a).

**Figure 6:**
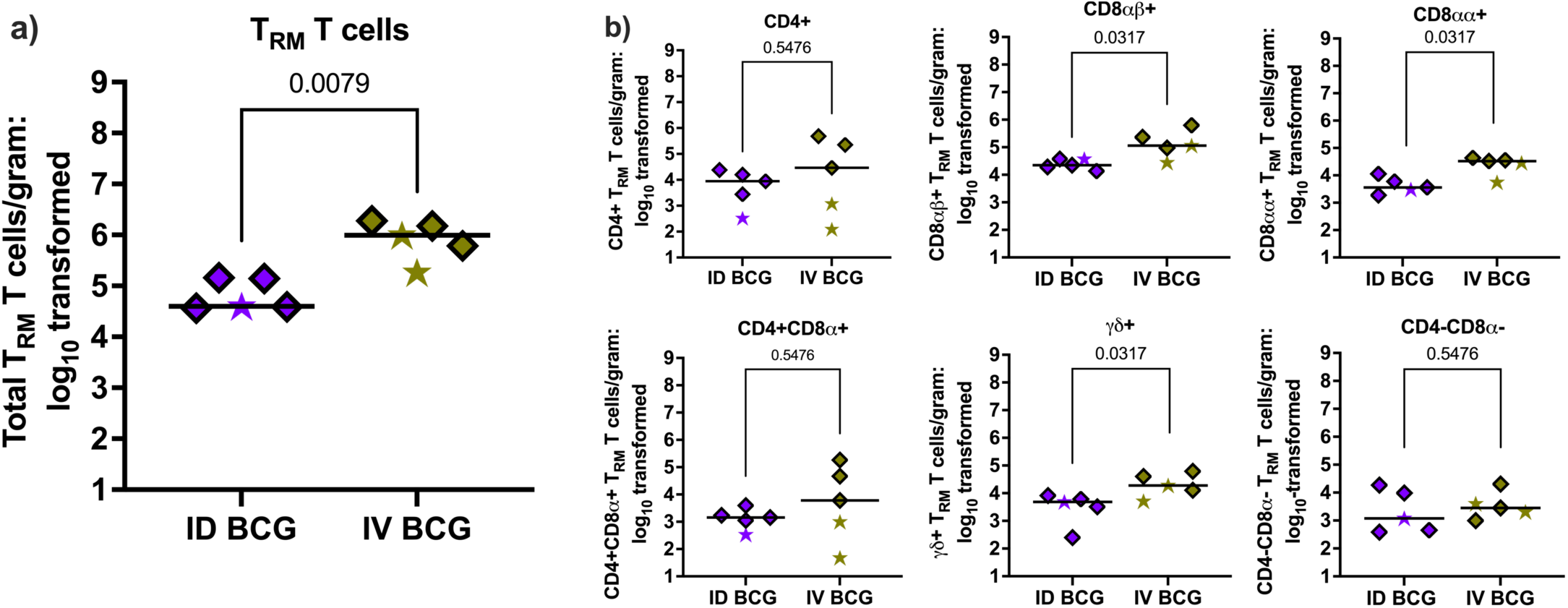
T_RM_ cell responses in lungs 20 weeks after BCG vaccination. Tissue-resident T cells (ivCD45-) are delineated from vascular-resident cells (ivCD45+). **a)** The number of tissue-resident T_RM_ cells in lung tissue harvested 20 weeks after IV or ID BCG vaccination. **b)** The number of T_RM_ cell subsets in lung tissue. SIV non-controllers are depicted by stars. Each symbol represents the mean value from 3 lung lobes per animal. Mann-Whitney p-values are reported.

### IV BCG induces higher CD4+ T cell cytokines but not CD8αβ+ T cells and NK cells cytotoxic effectors in lung tissue

We stimulated lung, thoracic lymph nodes, and spleen with WCL to assess production of cytokines associated with T1 (IFNγ, IL-2, TNF) and T17 (IFNγ, IL-2, IL-17, TNF) responses from CD4+ and CD8αβ+ T cells. The number of CD4+ T cells producing a combination of T1- and T17-associated cytokines in the lung were higher in animals vaccinated via IV BCG than ID BCG, but not in the thoracic lymph node (Figure 7a, 7b, and Supplemental Figure 2b, 2c). MCM with high SIV viremia had lower numbers of responding CD4+ T cells. CD8αβ+ T cells from all three sites produced some combination of T1 and T17 cytokines at similar levels in both IV- and ID-vaccinated animals (Figure 7a, 7b, and Supplemental Figure 2b, 2c).

**Figure 7:**
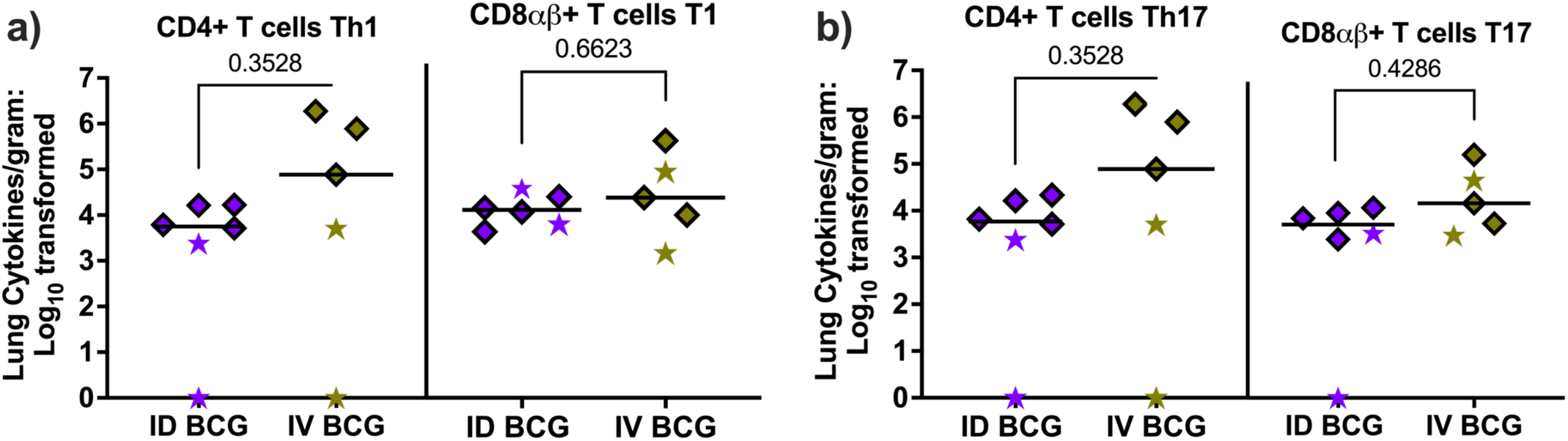
Cytokines responses in lungs 20 weeks after BCG vaccination. The number of CD4+ and CD8αβ+ T cells in the lung that make any **a)** Th1- and T1-(IFNγ, IL-2, TNF), and, **b)** Th17- and T17- (IFNγ, IL-2, IL-17, TNF) associated cytokines. SIV non-controllers are depicted by stars. Each symbol represents the mean value from 3 lung lobes per animal. Mann-Whitney p-values are reported.

We measured the number of CD8αβ+ T cells and NK cells producing granulysin, GzmB, GzmK, or perforin in lung, thoracic lymph nodes, and spleen. Our data revealed similar numbers of CD8αβ+ T cells and NK cells producing these cytotoxic effectors in lung and thoracic lymph nodes regardless of vaccination group (Supplemental Figure 2d-g). In the spleen, there were significantly more CD8αβ+ T cells producing GzmK and a trend towards more CD8ab+ T cells producing GzmB in IV- than ID-vaccinated animals (Data not shown), but NK cells producing cytotoxic effector show no significant difference in number in all the tissues (Supplemental Figure 2d-g).

There was no apparent effect of plasma SIV viremia on granulysin-, GzmB-, GzmK-, or perforin-producing CD8αβ+ T cells in the lung. However, spleens of IV BCG animals with higher plasma SIV viremia had the lowest GzmB-, GzmK-, or perforin-producing NK cells (Data not shown).

The transcription factors Foxp3, RORγT, and T-bet are associated with regulatory T cells, T17 cells, and T1 cells, respectively (44, 45). The number of CD4+ and CD8αβ+ T cells expressing Foxp3, RORγT, or T-bet did not differ in lung, thoracic lymph node, or spleen between animals vaccinated by the IV or ID route (Data not shown).

In summary, our data show that IV BCG induces robust and sustained T helper and cytotoxic T cell response in the airways of SIV+ MCM. Notably, even 5 months after vaccination, IV BCG-vaccinated animals had higher numbers of T_RM_ cells in the lungs compared to ID BCG-vaccinated animals.

## Discussion

PLWH are at increased risk from Mtb and a safe, effective TB vaccine for this population would be enormously beneficial. In SIV-naïve rhesus macaques, IV BCG vaccination provided superior protection from TB compared to ID BCG and correlated with mycobacterial-specific T cells in the airways and T_RM_ cells in lung tissue, mycobacterial-specific antibodies, and distinct transcriptional signatures (16, 20, 21, 46). We recently showed that IV BCG was also highly protective in MCM with preexisting SIV infection (19). Fully characterizing the immune responses induced by IV BCG in SIV+ macaques is important to further develop or learn from this vaccine strategy. However, it is difficult to differentiate immune responses in tissues induced by BCG from those induced by Mtb after vaccinated macaques are challenged with Mtb. Here, we investigated cellular and humoral immune responses to IV BCG in SIV-infected MCM in the blood, airways, and tissues without the confounding factor of Mtb challenge. We compared those to the responses induced by ID BCG in SIV-infected MCM as this route is used currently to vaccinate against TB. Our data demonstrate that in many ways the two vaccination routes induce similar immune responses in SIV+ macaques, but several features were unique to IV BCG-vaccinated animals, including some not investigated previously.

In our previous work using SIV-naïve rhesus macaques, IV BCG induced a robust T cell influx into airways, especially CD4+, CD8ab+, and nonconventional (Vg9+ gd and MAIT) T cells (16). We also showed a significant influx of these T cell subsets in SIV+ MCM after IV BCG vaccination (19). Here, we directly compared IV and ID BCG in SIV+ MCM and observed more CD4+ and CD8ab+ T cells in airways following IV vaccination. We also detected an increased influx of Vg9+ and Vg9-gd T cells into the airway following IV but not ID BCG as seen previously in SIV-naïve rhesus macaques (16, 47). We noted that more B and NK cells accumulated in the airways of the IV BCG-vaccinated, SIV-infected MCM. This is consistent with our previous study in SIV+ MCM (19) as well as our study in SIV-naive rhesus macaques (16). The latter study is particularly notable as the number of NK cells in the airways correlated with IV BCG-induced protection from Mtb (20). Thus, a key feature that differentiates IV BCG from ID BCG is a significant influx into airways of diverse lymphocyte populations which have all been shown to be associated with controlling Mtb in humans (48).

We also assessed the functionality of the cells that migrated to the airways in response to IV or ID vaccination. Both routes induced polyfunctional CD4+ T cells that produced T1- and T17-associated cytokines by 4 weeks. However, this response was more sustained in IV BCG-vaccinated animals, with significant elevation over 5 months. Polyfunctional T cells correlate with protection or improved outcome from Mtb in both humans and NHP (16, 20, 49–51), and we previously reported their induction following IV BCG vaccination in SIV+ MCM (19). In particular, Th1 and Th17 responses have been associated with protection from Mtb elicited by BCG delivered IV (20, 52) or intrabronchially (38). We showed here that CD8αβ+ T cell producing T1- and T17-related cytokines appeared in airways by 4 weeks after ID BCG but then waned over time. In contrast, T1- and T17-CD8αβ+ T cells were initially low following IV BCG but increased significantly by 18 weeks post-vaccination. Further, only IV BCG boosted the number of NK and CD8αβ+ T cells producing either GzmB or GzmK cytotoxic effector which had not been previously investigated. The protective efficacy of IV BCG may be related to the quality of the lymphocytes in the airways as well as the durability of those responses following vaccination.

Previous studies have shown that Mtb-specific antibody is associated with BCG-induced protection against Mtb in both SIV+ and SIV-naïve macaque (16, 19, 20, 52). Here, we demonstrated that IV BCG induces higher levels of mycobacterial-specific IgG and IgM in both the airways and plasma of SIV+ MCM than ID BCG. These antibodies were higher 4 weeks after vaccination than 18 weeks after vaccination and followed the kinetics of the B cells recovered from BAL. We previously reported increased mycobacterial-specific antibodies in both BAL and plasma from SIV+ MCM 12 weeks after IV BCG but did not assess other post-vaccination time points (19). These current results suggest that IV BCG recruits B cells to the airways and induces a robust IgG and IgM response a few weeks after vaccination which then subside. Since IV BCG-elicited IgM is associated with protection in rhesus macaques (21), it is likely that this humoral response remains primed to react quickly to Mtb challenge.

T_RM_ cells in the lung have been associated with protection from TB in mice (53) as well as in IV BCG-vaccinated macaques (16). In our previous study of IV BCG in SIV+ MCM (19), all animals underwent Mtb challenge and so we could not characterize vaccine-associated immune responses within tissues. The design of the current study did not include Mtb challenge and made it possible to evaluate T_RM_ cells within tissues in response to ID and IV BCG in SIV+ MCM. The use of intravascular anti-CD45 antibody (34) ensured that we analyzed tissue-resident, rather than circulating, immune cells. Our data revealed that even at 5 months post-vaccination, there were significantly more T_RM_ cells in lungs of SIV+ IV BCG-vaccinated macaques, compared to those vaccinated ID. These T_RM_ cells consisted of diverse T cell subsets, including CD4+, CD8αβ+, CD8αα+, CD8α+CD4+, and γδ+. In contrast, the number of T_RM_ cells in thoracic lymph nodes and spleen were similar between ID- and IV-vaccinated MCM, suggesting that IV BCG selectively promotes the accumulation and retention of T cells in lung tissue. Interestingly, the production of T1 and T17 cytokines, cytotoxic molecules, and transcription factors by these lung T cells did not differ between MCM vaccinated with ID or IV BCG, although the numbers of cells producing these effectors was increased in IV vaccinated animals. These data suggest that IV BCG confers protection, at least in part, by increasing the number of mycobacteria-specific T_RM_ cells in the lung rather than by inducing unique T cell responses. We previously showed a high numbers of polyfunctional CD4+ and CD8αβ+ T cell in lungs of SIV-naïve macaques 4 weeks after IV BCG vaccination (16). It is notable that, here, we evaluated lung-resident T cells 5 months post-BCG and observed higher numbers in IV BCG-vaccinated SIV+ animals which suggests that the increase in lung T_RM_ cells following IV BCG is quite durable. We speculate that IV BCG results in extensive T cell priming in lymph nodes and spleen and promotes their recruitment and retention in the lung, perhaps due to the persistence of BCG antigens following IV vaccination. This is supported by our previous finding in SIV-naïve macaques that viable BCG, albeit in low numbers, could be cultured from more organs 1 month after vaccination by the IV route compared to the ID route (16).

IV BCG induced a transient spike in SIV viremia following vaccination, as we reported previously (19). However, the current study was limited by a larger-than-expected number of animals with persistently high SIV viremia. Although the four non-controllers were distributed evenly between both experimental groups, it limited our ability to compare rigorously the immune responses in animals vaccinated by the IV and ID routes. These SIV non-controllers exhibited low numbers of CD4+ and CD4+CD8α+ T cells in blood and BAL as we seen previously (19, 25). These animals also had fewer CD4+ T_RM_ cells in lung tissue. Other cell types and antibody responses appeared unaffected by SIV viral loads. MCM with higher SIV loads were not protected from Mtb challenge (19), suggesting a detrimental effect of high viremia on the protective efficacy of IV BCG. Consequently, only three of five IV-vaccinated animals exhibited SIV control in our current study and, based on previous data (19), would have been likely to prevent TB.

In summary, the data presented here demonstrate clearly that BCG delivered to SIV+ macaques by the IV route leads to significantly higher numbers of lymphocyte populations in the airways, higher levels of mycobacterial-specific IgG and IgM, and more T_RM_ cells in the lung compared to ID vaccination. These findings inform potential mechanisms underlying the protection conferred by IV BCG in SIV+ MCM and provide valuable insights into vaccine design for PLWH.

## Acknowledgements

We are very grateful to the veterinary, laboratory, and analytical teams in the TB Research Group at the University of Pittsburgh for their dedicated work. We also thank Drs. Robert Seder and Patricia Darrah (VRC, NIAID, NIH) for helpful discussions.

