## supplemental Data for "Intravenous BCG induces a more potent airway and lung immune response than intradermal BCG in SIV-infected macaques^1^"

**Supplemental Figure 1**

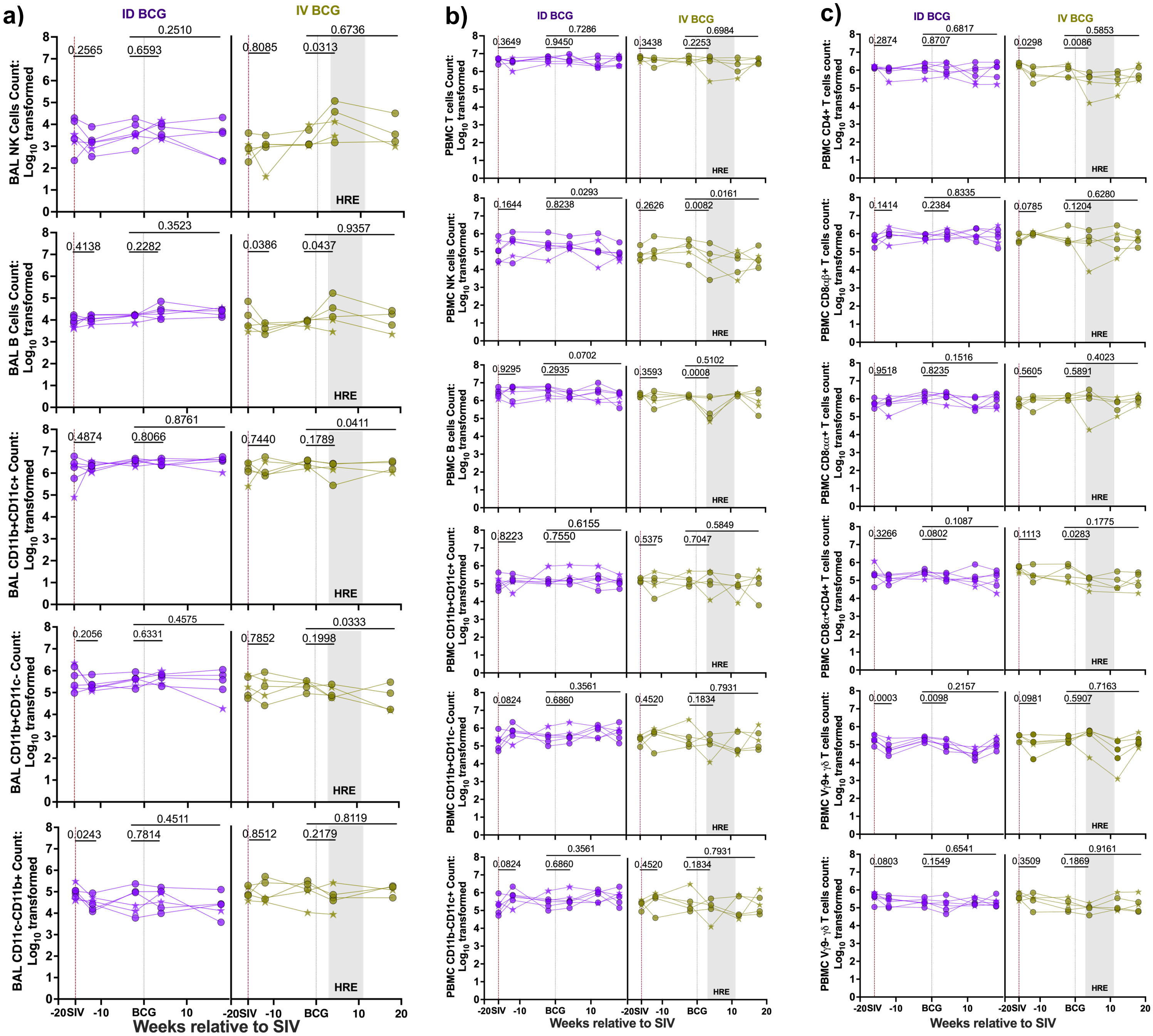

**Supplemental Figure 1: Leukocytes in blood and BAL and T cells in the blood in relation to SIV infection and BCG vaccination. a)**, and **b)** Number of total leukocytes in BAL and PBMC, **c)** number of total T cells in each PBMC. We employed mixed effects models, incorporating subject as a random variable, to evaluate variations across different time points and vaccine groups to evaluate changes that occurred between pre-SIV and 4 weeks post-SIV as well as pre-BCG and 4 weeks post-BCG. Mixed-effects model with uncorrected Fisher's LSD p-values reported above the time points of interest in each graph for changes that occur over time within a group.

**Supplemental Figure 2**

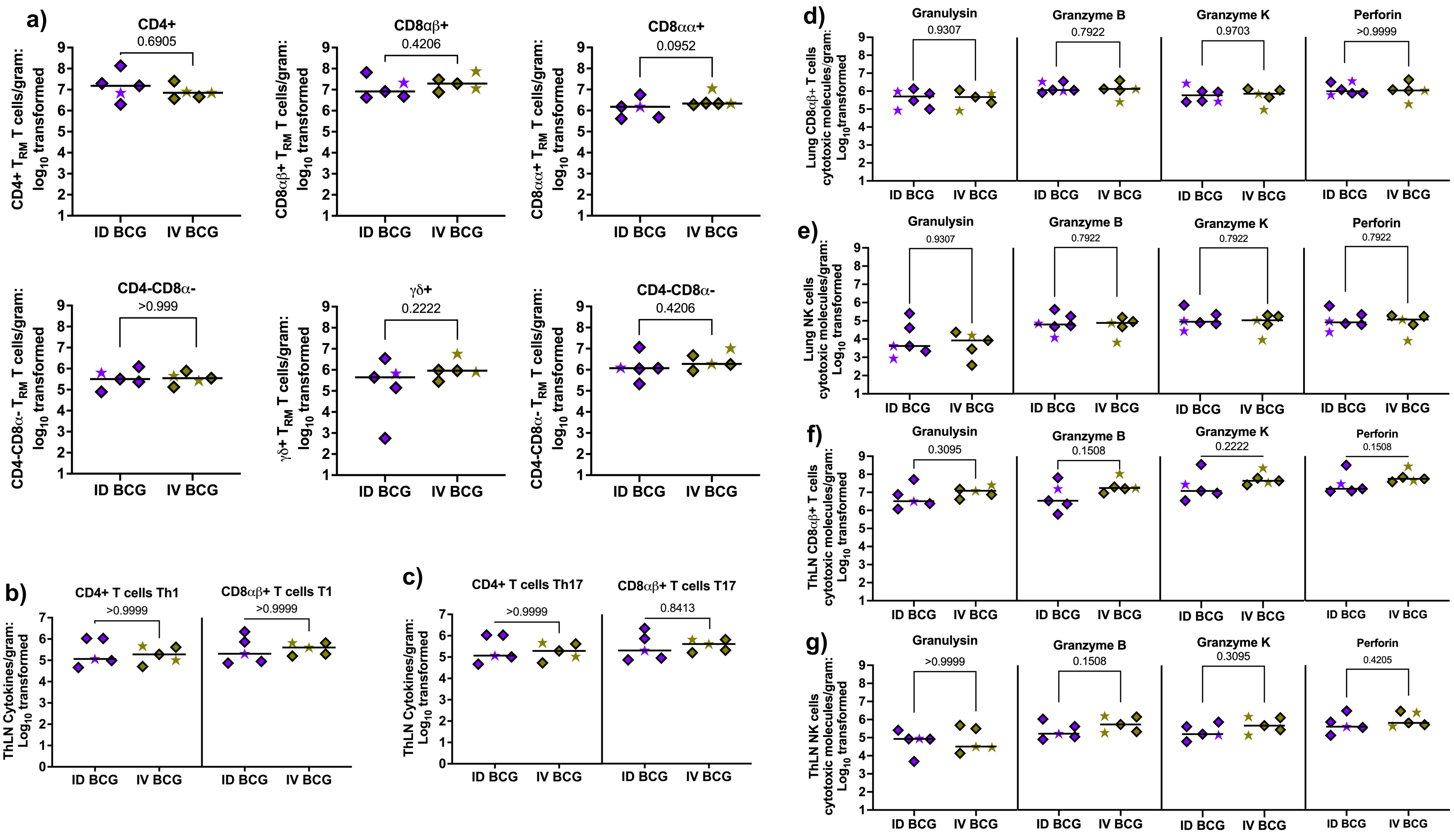


**Supplemental Figure 2: The quantification of T cells and Th1-, T1-, Th17- and T17-associated cytokines in thoracic LN and cytotoxic effector produced by CD8αβ+ T cell and NK cells in lung and ThLN 20 weeks post BCG. a)** T cells in the ThLN, **b)** and **c)** the number of cytokines producing CD4+ and CD8αβ+ T cells in ThLN. The number of granulysin, granzyme B, granzyme K, and perforin produced by CD8αβ+ T and NK cells in lung **d)** and **e)**, and ThLN **f)** and **g)**. The SIV virus controllers and non-controllers are shown in a diamond shape and star shape symbol, respectively. Each ThLN symbol represents a mean per MCM generated from 2 ThLN. Mann-Whitney p-values reported ID BCG (n=5) and IV BCG (n=5).

### Supplemental Figure 3

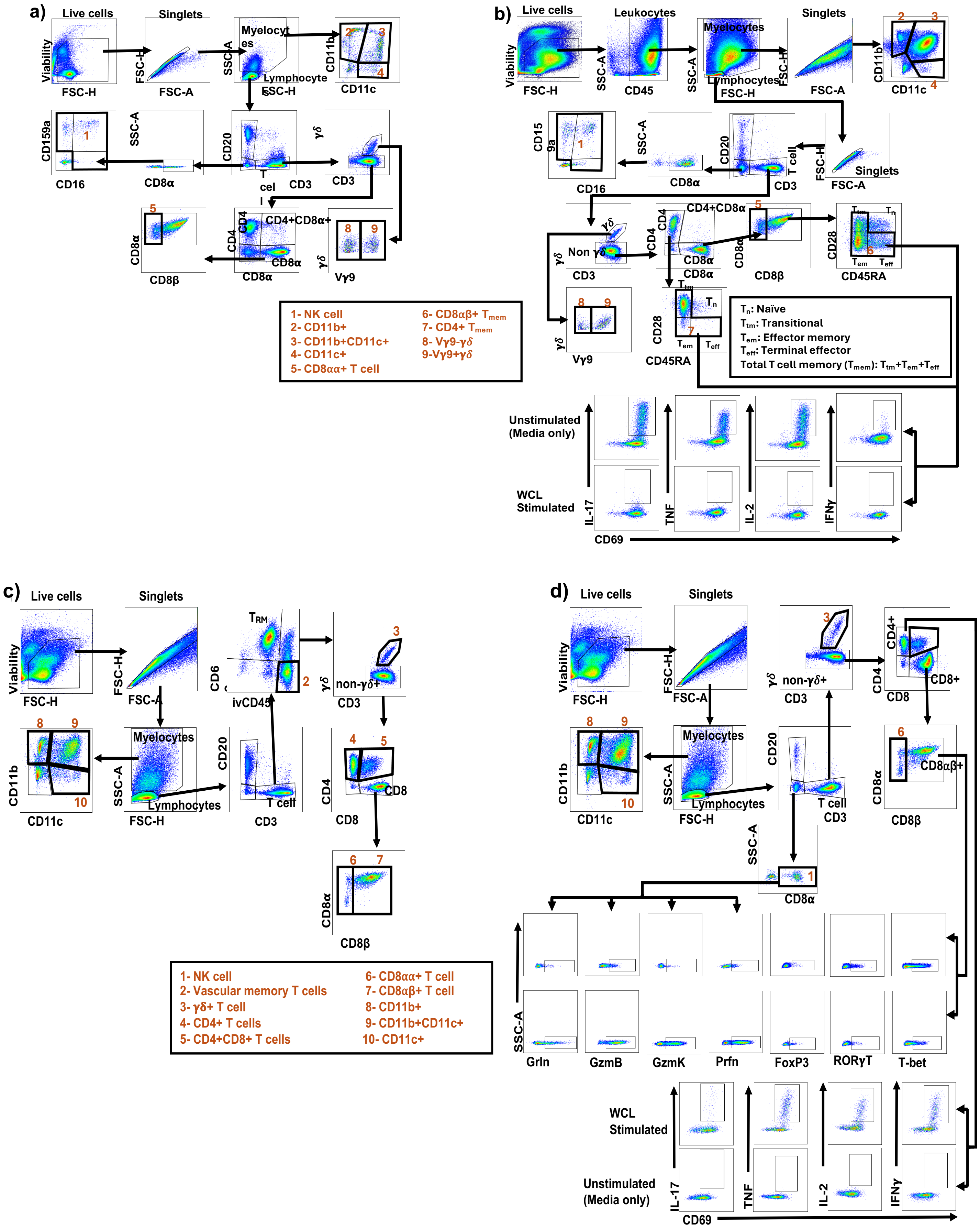

**Supplemental Figure 3. Gating schematic. Panel (a) PBMC and (b) BAL panel.** Cells were stained for flow cytometric analysis immediately after collection. Phenotypic cells (leukocytes and T cell subsets) were determined from unstimulated cells. Polyfunctional CD4+ and CD8+ T cell cytokines were determined after 6 hours of stimulation with WCL. BAL was first gated for live cells (Viability vs FSC-H), then gated leukocytes (SSC-A vs CD45), followed by myelocytes and lymphocytes gating (SSC-A vs FSC-H). Lymphocytes and myelocytes were gated for singlet, then myelocytes was characterized by CD11b vs CD11c, and lymphocyte was further gated into T cell, B cell and non-B_T cell. Non-B_T cell gated for CD8α then CD159a vs CD16 for NK cell. T cell was further differentiated into T cell subsets. Then CD4+ or CD8αβ+ T cell was gated to memory, namely Naïve T cells ($T_n$, CD45RA+CD28+); Transitional memory T cells ($T_{tm}$, CD45RA-CD28+); Effector memory T cells ($T_{em}$, CD45RA-CD28-); and Terminal effector T cells ($T_{eff}$, CD45RA+CD28-). Total memory ($T_{mem}$) was generated by OR gating of $T_{tm}$, $T_{em}$, and $T_{eff}$. B) CD4+ and CD8αβ+ T cell memory ($T_{mem}$) response gating for IFNγ, TNF, IL-2, and IL-17A in the presence of no stimulation (media only) or PPD. Similar gating schematic was on both C4+ and CD8αβ+ T cells $T_{mem}$ for cytokines. **Panel c) and d) show tissue gating schematic,** tissue samples collected at necropsy were freshly homogenized and passed through a cell strainer to obtain single cells for flow staining. **a)** Unstimulated tissue-resident T ($T_{RM}$) cell was delineated from blood-derived T cell. It was measured in the tissue by injecting MCM with a fluorochrome-labeled antiCD45 (ivCD45) before necropsy. CD69 was used as the tissue-resident maker, therefore, the $T_{RM}$ cells was labeled ivCD45-CD69+. **b)** Gate from single cell suspension obtained freshly from necropsy were split into stimulated (using WCL) and unstimulated and stained for flow cytometry to quantify T cell subsets and polyfunctional cytokines producing CD4+ and CD8αβ+ T cells. IFNγ, TNF, IL-2, and IL-17 were gated individually, then CD4+ T cell producing Th1 (IFNγ, TNF, and IL-2) and Th17 (IFNγ, TNF, IL-2, and IL-17A) cytokines, and CD8αβ+ T cell producing T1 (IFNγ, TNF, and IL-2) and T17 (IFNγ, TNF, IL-2, and IL-17A) cytokines were generated by OR gate. Grln: Granulysin, GzmB: Granzyme B, GzmK: Granzyme K and Prfn: Perforin

**Supplemental Table 1: List of antibodies used in various flow panel**

| Marker | Fluorophores | Clone | Panel | Stain target | Company |
|---|---|---|---|---|---|
| | | | **Surface stain** | | |
| CD45 | BV480 | D058-1283 | PBMC phenotype, BAL phenotype | Surface | BD Biosciences |
| ivCD45 | AF488 | MB4-6D6 | Tissue | Surface | Miltenyi Biotec |
| CD3 | APC-Cy7 | SP34-2 | PBMC phenotype, BAL phenotype, tissue | Surface | BD Biosciences |
| Pan_δγ | PE | 5A6.E9 | PBMC phenotype, BAL phenotype, tissue | Surface | Thermofisher |
| Vγ9 | FITC | 7A5 | PBMC phenotype, BAL phenotype | Surface | Thermofisher |
| CD4 | BV750 | L200 | PBMC phenotype, BAL phenotype | Surface | BD Biosciences |
| CD4 | BUV563 | SK3 | Tissue | Surface | BD Biosciences |
| CD8α | RPA-T8 | BUV496 | PBMC phenotype, BAL phenotype | Surface | BD Biosciences |
| CD8α | RPA-T8 | BUV395 | Tissue | Surface | BD Biosciences |
| CD8β | BUV737 | 2ST8.5H7 | PBMC phenotype, BAL phenotype, tissue | Surface | BD Biosciences |
| CD45RA | BUV395 | 5H9 | PBMC phenotype, BAL phenotype | Surface | BD Biosciences |
| CD28 | BV605 | CD28.2 | PBMC phenotype, BAL phenotype | Surface | BioLegend |
| CD20 | PE-CY5 | 2H7 | PBMC phenotype, BAL phenotype | Surface | BioLegend |
| CD20 | BV570 | 2H7 | Tissue | Surface | BioLegend |
| CD159a | PE-CY7 | Z199 | PBMC phenotype, BAL phenotype | Surface | Beckman Coulter |
| CD16 | BV570 | 3G8 | PBMC phenotype, BAL phenotype | Surface | BioLegend |
| CD11b | BUV563 | ICRF44 | PBMC phenotype, BAL phenotype | Surface | BD Biosciences |
| CD11b | BV510 | ICRF44 | Tissue | Surface | BioLegend |
| CD11c | BV421 | 3.9 | PBMC phenotype, BAL phenotype | Surface | BioLegend |
| CD11c | AF700 | 3.9 | Tissue | surface | Biolegend |
| CD69 | BV711 | FN50 | Tissue | Surface | BioLegend |
| Live/Dead | Zombie NID | | PBMC phenotype, BAL phenotype | surface | BioLegend |
| Live/Dead | Live/Dead Blue | | Tissue | Surface | Thermofisher |
| | | | **Intracellular Stain** | | |
| CD69 | ECD | TP1.55.3 | PBMC phenotype, BAL phenotype | Intracellular | Beckman Coulter |

| | | | | | |
|---|---|---|---|---|---|
| CD69 | BV711 | FN50 | Tissue | Intracellular | BioLegend |
| Granulysin | AF647 | eBioDH2 (DH2) | Tissue | Intracellular | Thermofisher |
| Granzyme B | V450 | GB11 | PBMC phenotype, BAL phenotype, tissue | Intracellular | BD Biosciences |
| Granzyme K | AF647 | G3H69 | PBMC phenotype, BAL phenotype | Intracellular | BD Biosciences |
| Granzyme K | G3H69 | PerCP-eFluor 710 | Tissue | Intracellular | Invitrogen |
| IL-2 | BV785 | MQ1-17H12 | PBMC phenotype, BAL phenotype | Intracellular | BioLegend |
| IL-2 | BV750 | MQ1-17H12 | Tissue | Intracellular | BD Biosciences |
| IL-17 | BV510 | BL168 | PBMC phenotype, BAL phenotype | Intracellular | BioLegend |
| IL-17 | PE-Cy7 | BL168 | Tissue | | BioLegend |
| IFNγ | APC | B27 | PBMC phenotype, BAL phenotype | Intracellular | Biolegend |
| IFNγ | BV785 | 4S.B3 | Tssue | Intracellular | BioLegend |
| TNFα | BV650 | MAb11 | PBMC phenotype, BAL phenotype, tissue | Intracellular | BioLegend |
| Perforin | PE-Cy5 | PF-80/164 | Tissue | | Mabtech |
| **Intranuclear stain** | | | | | |
| FoxP3 | V450 | 259D/C7 | Tissue | Intranuclear | BD Biosciences |
| RORγT | APC | AFKJS-9 | Tissue | | Thermofisher |
| T-bet | PE-Cy5 | eBio4B10 (4B10) | Tissue | Intranuclear | Thermofisher |
